# Metabolic Architecture of the Deep Ocean Microbiome

**DOI:** 10.1101/635680

**Authors:** Silvia G. Acinas, Pablo Sánchez, Guillem Salazar, Francisco M. Cornejo-Castillo, Marta Sebastián, Ramiro Logares, Shinichi Sunagawa, Pascal Hingamp, Hiroyuki Ogata, Gipsi Lima-Mendez, Simon Roux, José M. González, Jesús M. Arrieta, Intikhab S. Alam, Allan Kamau, Chris Bowler, Jeroen Raes, Stéphane Pesant, Peer Bork, Susana Agustí, Takashi Gojobori, Vladimir Bajic, Dolors Vaqué, Matthew B. Sullivan, Carlos Pedrós-Alió, Ramon Massana, Carlos M. Duarte, Josep M. Gasol

## Abstract

The deep sea, the largest compartment of the ocean, is an essential component of the Earth system, but the functional exploration of its microbial communities lags far behind that of other marine realms. Here we analyze 58 bathypelagic microbial metagenomes from the Atlantic, Indian, and Pacific Oceans in an unprecedented sampling effort from the Malaspina Global Expedition, to resolve the metabolic architecture of the deep ocean microbiome. The Malaspina Deep-Sea Gene Collection, 71% of which consists of novel genes, reveals a strong dichotomy between the functional traits of free-living and particle-attached microorganisms, and shows relatively patchy composition challenging the paradigm of a uniform dark ocean ecosystem. Metagenome Assembled Genomes uncovered 11 potential new phyla, establishing references for deep ocean microbial taxa, and revealed mixotrophy to be a widespread trophic strategy in the deep ocean. These results expand our understanding of the functional diversity, metabolic versatility, and carbon cycling in the largest ecosystem on Earth.

**One Sentence Summary:** A whole community genomic survey of the deep microbiome sheds light on the microbial and functional diversity of the dark ocean.

## Main Text

Most of the ocean’s life is isolated from our planet’s primary energy source, the sun light. Besides living in permanent darkness, deep ocean organisms have to cope with high pressure and low temperature, which makes life in this ecosystem challenging. This fascinating habitat represents one of the largest biomes on Earth, mostly occupied by bacteria and archaea that play a pivotal role in biochemical cycles on a planetary scale (Cho and Azam 1988, Herndl and Reinthaler 2013, Bar-On *et al*., 2018). Microbial metabolisms in the deep ocean have been assumed to be primarily heterotrophic relaying on organic matter exported from the sunlit layer through sinking particles, which support the bulk (90%) of the respiratory carbon demand of the dark ocean (Arístegui *et al*., 2009). However, the high respiratory activity measured in the dark ocean is difficult to reconcile with the rates of supply of organic carbon produced in the photic layer (del Giorgio and Duarte 2002, Arístegui *et al*., 2003, 2005, 2009, Herndl and Reinthalerl 2013), suggesting the need for significant autotrophic production in the aphotic ocean. Chemolithoautotrophy has been considered as a possible pathway supporting this high dark ocean respiratory activity (Herndl *et al*., 2005, Reintahler *et al*., 2010, Herndl and Reinthaler 2013). Experimental rate measurements and bulk biogeochemical estimates agree with the findings of potential for chemolithoautotrophy in ubiquitous bacterial and archaeal lineages using single cell genomics (Swan *et al*., 2011, Pachiadaki *et al*., 2017). These results suggest that deep-sea chemolithoautotrophy may not be a marginal contributor to deep-sea metabolism and may play a greater role in the global ocean carbon cycle than previously thought. Whereas inorganic carbon fixation is energetically costly (Hügler and Sievert, 2011), mixotrophy, i.e. simultaneously carrying out chemolithoautotrophic inorganic carbon fixation and heterotrophy (Sorokin, 2003, Dick *et al*., 2008, Alonso-Sáez *et al*., 2010), may constitute a cost effective strategy for microorganisms to persist in the dark ocean.

Particles represent important hot-spots of microbial activity fueling the dark ocean food web (Herndl and Reinthaler 2013). These particles include fast-sinking, traveling through the water column in a few weeks (Turner 2002; Ploug *et al*., 2008; Agustí *et al*., 2015), as well as buoyant or slow-sinking organic particles, which remain suspended in the deep ocean over annual times scales (Herndl and Reinthaler 2013). The diversity and biogeography of bathypelagic prokaryotic communities was only recently described at a global scale, showing that the free-living (FL) and particle-attached (PA) microbial communities differ greatly in taxonomic composition and appear to be structured by different ecological drivers (Salazar *et al*., 2016a, Mestre et al. 2018). The lifestyle dichotomy between free-living and particle attached prokaryotes was shown to be a phylogenetic conserved trait of deep ocean microorganisms (Salazar *et al*., 2016b); however, the differences in the functional capacities of these two groups of microorganisms remain largely unknow. A global-level understanding of the ecology and metabolic processes of deep-sea microorganisms in this heterogeneous environment, comparable to that already available for microorganisms in the upper ocean (DeLong *et al*., 2006; Rusch *et al*., 2007; Sunagawa *et al*., 2015, Brum *et al*., 2015, Roux *et al*., 2016, Louca *et al*., 2016, Carradec *et al*., 2018) is still missing.

The Malaspina 2010 Circumnavigation Expedition addressed these gaps by carrying out a global survey of bathypelagic microbes in the tropical and subtropical oceans (Duarte 2015). We use metagenomics to explore and map the main biochemical processes in the deep ocean and we do so by generating the Malaspina Deep-Sea Gene Collection, which we offer here as valuable deep-ocean microbial community genomic dataset, and by reconstructing metagenomic assembled genomes (MAGs) to explore the metabolic capacities of the deep-sea microbiome. In particular, we report and analyze 58 deep-sea microbial metagenomes sampled between 35°N and 40°S from 32 stations (St) in the N and S Pacific and Atlantic Oceans, the Indian Ocean, and the South Australian Bight (**Fig. 1A**) with an average depth of 3,731 m. Two different plankton size fractions were analyzed in each station representing the free-living (FL, 0.2–0.8 µm) and particle-attached (PA, 0.8–20 µm) microbial communities.

**Figure 1.**
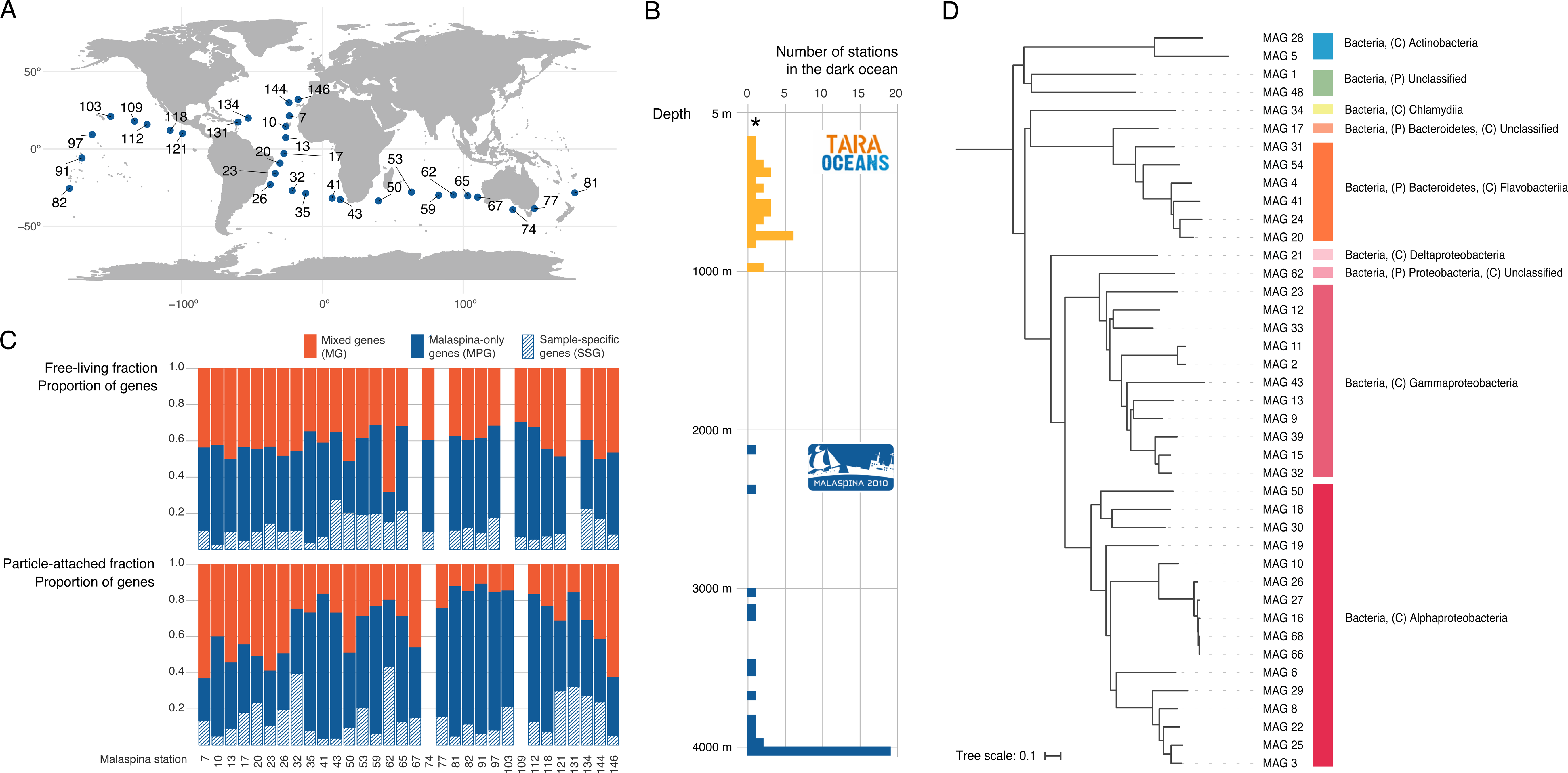
Malaspina Deep Ocean Genetic Resources. A) Malaspina 2010 expedition station map showing the location of the 32 stations sampled along the circumnavigation used in this study. B) Representation of the sampling depth and metagenomics dataset generated by the Tara Oceans and Malaspina 2010 Circumnavigation Expeditions. The histogram plot displays the number of stations sampled in the dark ocean in the Tara Oceans (orange) and Malaspina 2010 (blue) expeditions. The Malaspina Deep-Sea Gene Collection was generated from the integration of 58 metagenomic bathypelagic samples. The asterisk in the histogram indicates that samples collected in the photic layer in Tara Oceans are not included in the plot. C) Analyses of the integrated gene catalogue that results from the Tara Ocean (OM-RGC) and Malaspina Deep-Sea Gene Collection. Relative abundance of unique genes that appear only in Malaspina (MPCL) (solid blue), Mixed genes (MCL) that include genes from both catalogues (red) and the Malaspina Sample-Specific genes (SSCL) in shaded blue for both the Free-Living (FL; 0.2-0.8 µm) and Particle-Attached (PA; 0.8-20 µm) fractions. D) Phylogenetic analyses of 41 out of 76 metagenomic assembled genomes (MAGs) obtained from co-assembling 58 bathypelagic ocean metagenomes. A total of 41 MAGs were displayed at the maximum taxonomic possible resolution (P= phylum, C=class), specifically those that taxonomy agreed in the two methods applied: i) the taxonomy between the one estimated by checkM and 2) the one assigned by homology search against UniRef100 of all coding genes from each MAG (least common ancestor).

### The Malaspina Deep-Sea Gene Collection

Shotgun sequencing (paired-end 2×150 bp) using the Illumina HiSeq2000 platform generated a total of 195 gigabases (Gb) (6.49×10^8^ read pairs) of data with an average of 3.36 Gb per sample (**Table S1**). The resulting Malaspina Deep-Sea Gene Collection (MDSGC) built from these 58 bathypelagic metagenomes contained a total number of 4.03 million (M) genes predicted in the assembled contigs (**Table S1**). This dataset from the bathypelagic ocean offers a unique gene repertory complementary to the mesopelagic metagenomes reported by *Tara* Oceans (**Fig. 1B**). The MDSGC, once clustered at 95% sequence identity to remove gene redundancy, yielded 1.12 M non-redundant unique gene clusters (referred hereafter as *genes*), that were then analyzed and integrated into the published *Tara* Oceans OM-RGC.v1 (see methods, **Table S1**) (**Fig.1C**). Remarkably, 71% of the non-redundant unique gene sequences found in the bathypelagic metagenome had not been detected in the global surveys of the upper ocean (Sunagawa *et al*., 2015), which is surprising given the role of sinking particles in delivering epipelagic microbes to the deep-sea (Mestre *et al*., 2018). These 71% of the genes unique to the Malaspina deep-sea gene collection represented a total of 657,469 non-redundant Malaspina-exclusive genes (MPG) (**Fig. 1C**). On average (± SD) 62.9 ± 13.7% of the predicted exclusive genes in each sample (MPG) corresponded to new genes, which highlights the remarkably novel gene content of the bathypelagic microbiome (**Fig. 1C, Table S1**). On average, we detected 14.1 ± 8.9% sample-specific genes (SSG) (**Fig. 1C, Table S1**) in the Malaspina samples. Station St62, in the Indian Ocean, showed the highest fraction of sample-specific genes with 43.3% of the total. This sample, together with other 4 stations with more than 30% sample-specific genes located in the Brazil (St32), North Atlantic American (St131) and Guatemala basins (St121) were all from the particle-attached size fraction and associated with circumpolar deep water (CDW) and North Atlantic Deep Water (NADW) as the dominant water masses (**Table S1).**

The taxonomy of the microbes (i.e. small eukaryotes, prokaryotes, and viruses) in the bathypelagic samples was evaluated using different marker genes extracted from metagenomes (**Fig. S1, see methods**). The protistan, bacterial, and archaeal diversity patterns of the deep ocean confirmed previous results based on 18S (Pernice *et al*., 2016) and 16S (Salazar *et al*., 2015a) PCR amplicons on the same samples. For protists, the main difference with the photic layers was the relevance of Excavates along with the remarkable importance of fungal taxa in the deep ocean (**Fig. S1A**). For bacteria and archaea, *Thaumarchaeota*, accounted for 21% of all sequences, dominating in the FL fraction (**Fig. S1B**) and as expected, a marginal presence of cyanobacteria and higher abundances of Gammaproteobacteria (27.3%) and Actinobacteria (17.3%) in the deep ocean compared to the dominance of Alphaproteobacteria in the photic layers (Sunagawa *et al*., 2015). Identification of nucleo-cytoplasmic large DNA viruses (NCLDVs) revealed the presence of NCLDVs in all sampled size fractions across ocean basins (**Fig. S1C**). Megaviridae, whose known hosts are either mixotrophic or heterotrophic unicellular eukaryotes, were found to be the dominant group of NCLDVs in deep oceans (76% and 63% in the FL and PA fractions, respectively), which contrasts with the lower proportion (36%) of Megaviridae in the sunlit ocean (Hingamp *et al*., 2013). Finally, the marker gene *terL* covered the three *Caudovirales* families (*Myoviridae, Podoviridae, Siphoviridae*), which are the dominant type of viruses infecting bacteria and archaea in mesopelagic environments (Brum *et al*., 2015). The 485 *terL* genes from the Malaspina deep-sea gene collection were mostly novel (**Fig. S1D**).

#### Functional architecture of the deep ocean microbiome: free living vs. particle-attached assemblages

We analyzed the Malaspina Deep-Sea Gene Collection to evaluate the differences in functional structure between bacteria and archaea of the particle-attached size fraction and those living in the surrounding water. This is important, because functional analyses of marine FL and PA microorganisms are limited to a few studies at the local scale in the photic ocean (Allen *et al*., 2012; López-Pérez *et al*., 2015), coastal ecosystems (Smith *et al*., 2013) or the oxygen minimum zone (OMZ) off the coast of Chile (Ganesh *et al*., 2014). All these studies showed contrasting gene repertories for FL and PA microbial communities. Our results also identified two main functional groups of samples corresponding to particle-attached and free-living communities (**Fig. 2**). This pattern was coherent regardless of the functional classifications used based on Kyoto Encyclopedia of Genes and Genomes Orthology (KOs) (**Fig. 2**), clusters of orthologous groups (COG), protein families (Pfam) or Enzyme Commission numbers (EC) (**Fig. S2**). Our results showed that particle-attached and free-living microbial communities represent different functional lifestyles with different metabolisms and functional traits.

**Figure 2.**
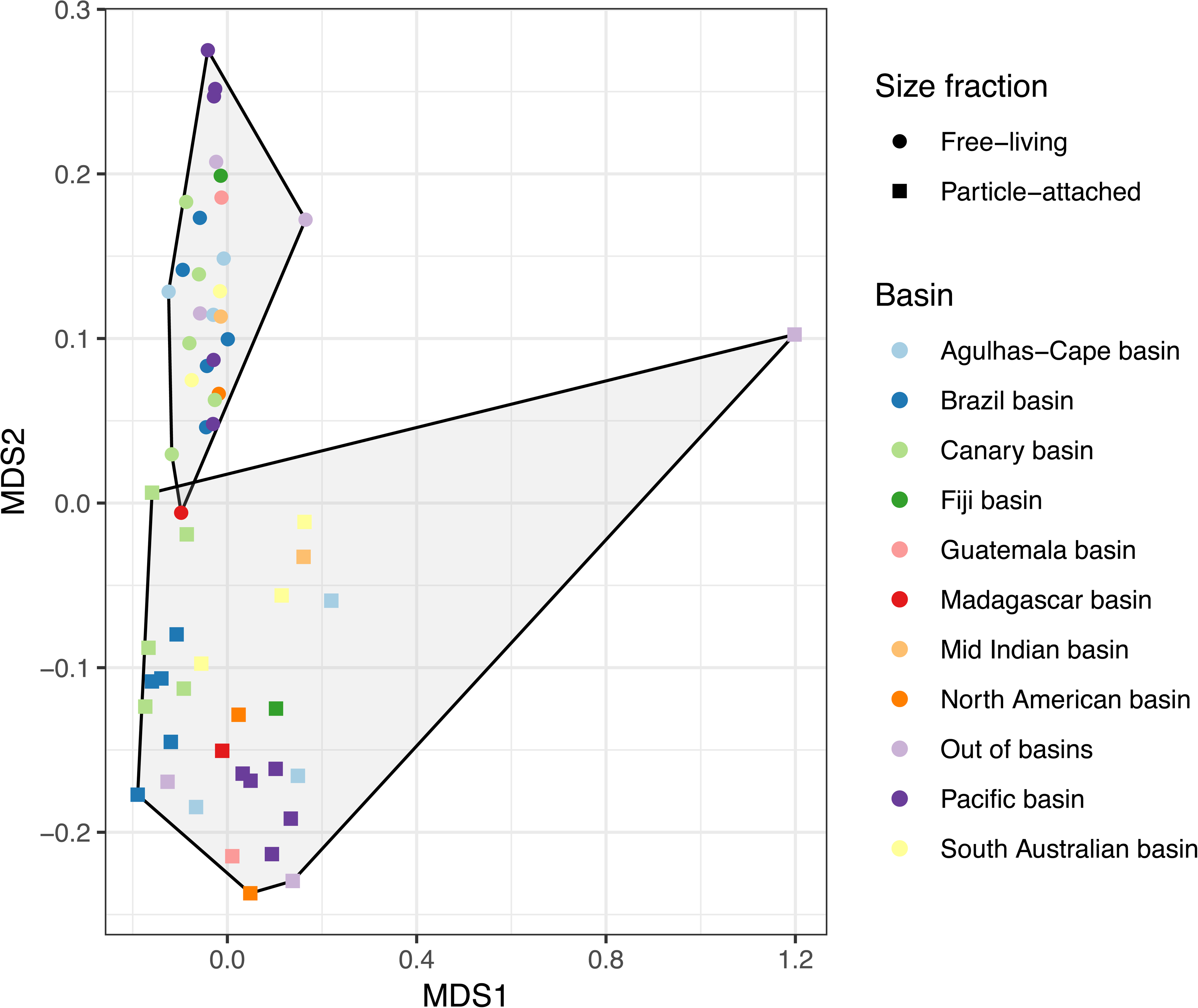
Functional community structure of the bathypelagic microbial communities. A) Non-metric multidimensional scaling (NMDS) of the microbial communities based on the functional compositional similarity (Bray-Curtis distances) among the 58 samples in the dataset, based on clusters of KEGG orthologous groups (KOs). Size-fraction is coded by symbol (squares, particle-attached and circles, free-living prokaryotes) and the main deep-oceanic basins by color codes (see legends).

To explore the ubiquity of the main biogeochemical cycling metabolisms present in the deep ocean involving inorganic carbon fixation, nitrogen, sulfur, methane, and hydrogen related metabolisms (**Fig. 3)** and the potential differences in the FL and PA microbial communities (**Fig. 4**), a selection of key marker genes (69 KOs, see **Table S2**) were analyzed and normalized by *recA*, a single copy gene as a proxy for copy number per cell (**Fig. 3, Table S3**). We detected a total of 40 KOs in the Malaspina Deep-Sea Gene Collection related to deep ocean biogeochemical processes, including the key genes of four different inorganic CO_2_ fixation pathways. Two of them, the Calvin cycle and the 3-hydroxypropionate bi-cycle (3-HP) were widely distributed in our dataset while, the archaeal 3 hydroxypropionate-4 hydroxybutyrate cycle (K15039), and the reverse TCA cycle (rTCA; K15230/ K15231) displayed a narrow distribution restricted to a single station (**Fig. 3).** First, the key enzyme of the Calvin Cycle, the ribulose-bisphosphate carboxylase (rubisco, *rbcL*, K01601) was present in 48% of the samples (**Table S3**). The *rbcL* gene was present on average in 1.3% of the cells, but peaked up to 12% in the particle-attached fraction of St67 (MP1241; South Australian Bight, Indian Ocean) and St134 (MP2633, North American basin, Atlantic Ocean) (**Fig. 3).** Also relevant was the presence of the 3-hydroxypropionate bicycle in which two key enzymes such as the malonyl-CoA reductase (K14468: *mcr*) and malyl-CoA lyase (K08691: *mcl*) (Zarzycki *et al*., 2009) occurred in 1.7 % and 52% of the samples respectively and in 2% of the cells and with a maximum peak at the station St82 (MP1493) at the Pacific Ocean in 22% of the cells (**Fig 3**). The other two enzymes detected from the 3-hydroxypropionate bi-cycle pathway (K09709: *meh* and K14470: *mct*) were also found in at least half of the samples. The 3-hydroxypropionate bi-cycle pathway, which uses HCO_3_^−^ as inorganic carbon species, was originally associated with the green nonsulfur bacterium *Chloroflexus aurantiacus* and with other members of *Chloroflexaceae* (Strauss and Fuchs 1993, Zarzycki *et al*., 2009). However, individual genes of the pathway have been detected in various strains of Alpha- and Gammaproteobacteria (Zarzycki *et al*., 2009) and recently in deep ocean SAR202 clade genomes (Landry *et al*., 2017, Mehrshad *et al*., 2018). These results not only reflect a ubiquitous role for some inorganic carbon fixation pathways in the deep ocean, highlighting the wide distribution of the Calvin Cycle and the 3-Hydroxypropionate bicycle in both PA and FL microbial communities (**Fig. 4**), but also indicate a relevant degree of patchiness in their appearance in the deep ocean.

**Figure 3.**
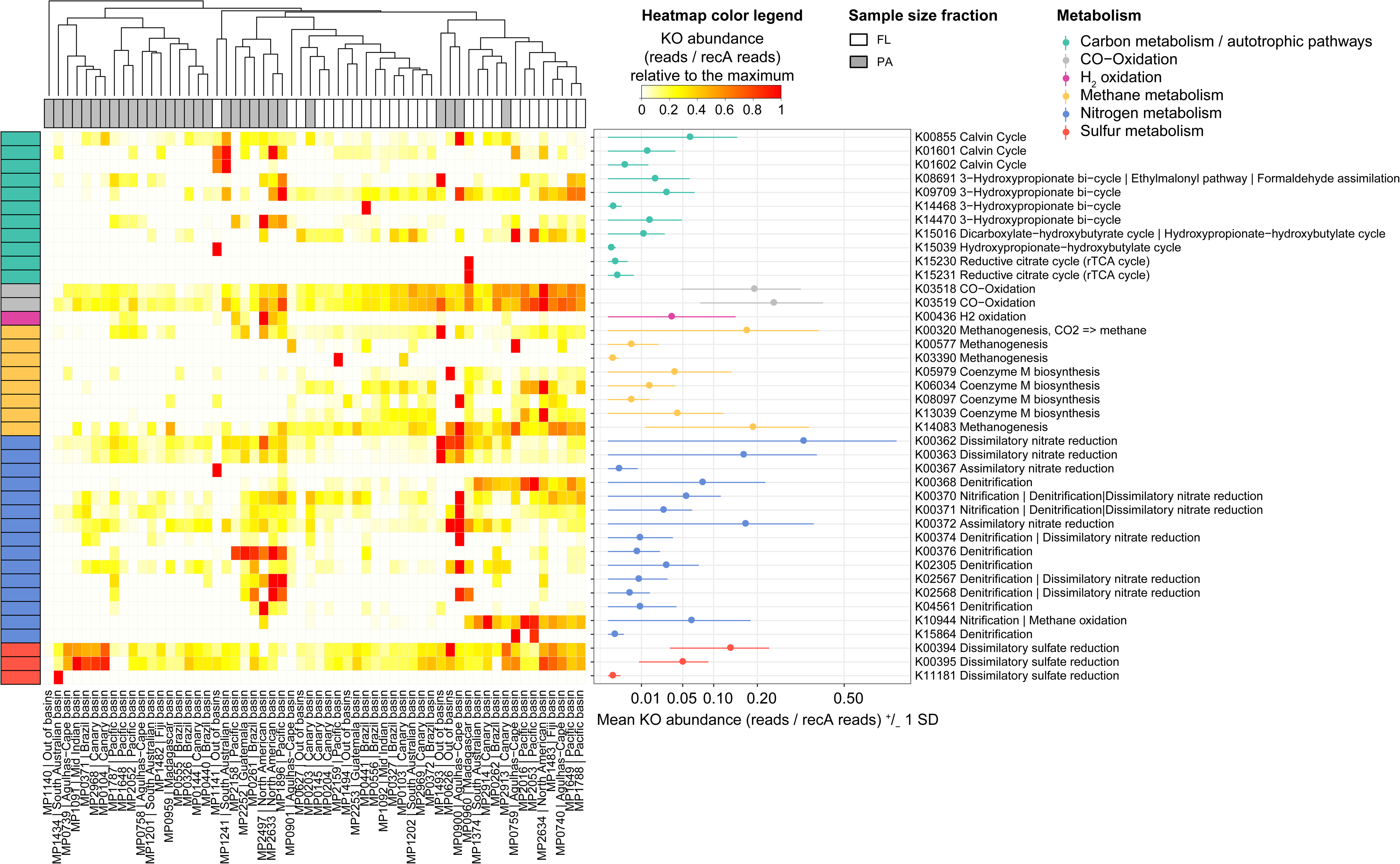
Heat-map of selected markers genes for different metabolic pathways across the 58 metagenomes. A total of 40 marker genes (KOs, Y axis) indicative of different metabolic processes (Table S4) detected in the Malaspina samples (X axis). KO abundance was normalized by *recA* single copy gene as a proxy for copy number per cell. The general metabolism assignation is color-coded (see legend in the upper right) and the KEGG module(s) assignation used in the KO label is also indicated. The relative abundance across samples for each KO is shown in the heatmap. The mean (± 1 s.d.) untransformed abundance of each KO across all samples (reads / *recA* reads) is presented in the right panel.

**Figure 4.**
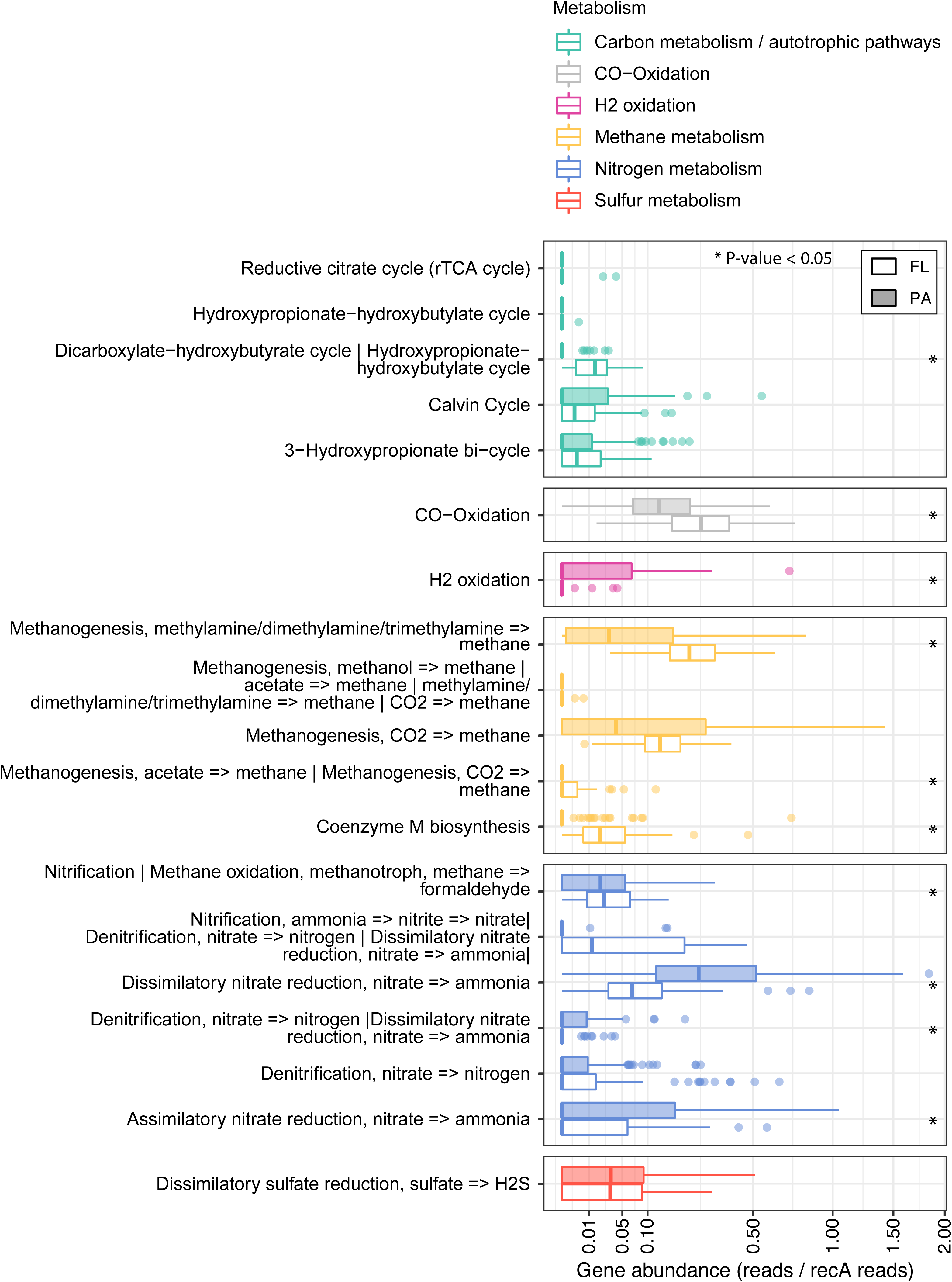
Comparison of metabolic pathways in the free-living and particle-attached dark ocean microbiomes. The normalized gene abundance (reads / *recA* reads) of 40 marker genes indicative of different autotrophic carbon fixation pathways, nitrogen, sulfur and methane metabolisms in the Malaspina Deep-Sea Gene Collection. Gene abundance are summarized at the KEGG module level. Wilcoxon tests were done to test for significant differences between the particle-attached (PA) and free-living (FL) assemblages and significant (p-value < 0.05) differences are labeled with asterisks. Free-living (empty boxes) and particle-attached (filled boxes) bathypelagic microbial communities are shown next to each other.

Nitrification through ammonia (Wuchter *et al*., 2006; Alonso-Saez *et al*., 2012) and nitrite oxidation (Pachiadaki *et al*., 2017) has been postulated as the main source of energy for carbon fixation in the dark ocean. The corresponding gene marker (methane/ammonia monooxygenase subunit A, K10944; *pmoA*-*amoA*), although not only associated to ammonia oxidation, was detected in 36% of our samples and 6% of the microbial cells and linked to free-living microorganisms (Wilcoxon test p-value < 0.005) (**Fig. 4).** The K00370/K00371 (nitrate reductase/nitrite oxidoreductase) could be related to nitrite oxidation and was found in 81% of the samples (**Fig. 3**) and in about 4% of the cells (**Fig. 4**). However, other relevant nitrification pathways involving the hydroxylamine dehydrogenase (hao; K10535) or the key enzyme for the anaerobic ammonia oxidizers (anammox bacteria, K20932/K20935), normally associated within the oxygen minimum zones in the mesopelagic ocean, were absent from our bathypelagic dataset, reflecting different biogeochemical processes occurring in the mesopelagic and the bathypelagic oceans.

H_2_-oxidation and CO-oxidation were explored as alternative energy sources in the deep ocean. The H_2_-oxidation (K00436: hoxH gene) is usually associated to hydrothermal vents (Anantharaman *et al*., 2012) or subsurface microbial communities (Brazelton *et al*., 2012) but we found it in 24% of the samples and in (**Fig. 3**), mainly associated to particle-attached microorganisms (Wilcoxon test p-value < 0.005) (**Fig. 4**). These results expand the ecological niches of H_2_ oxidizers in the bathypelagic ocean, probably associated to particles providing anoxic microenvironments where H_2_ production by fermentation is favorable. The oxidation of carbon monoxide by aerobic carboxydotrophs is catalyzed by carbon monoxide dehydrogenase (CODH; cox genes) (Ragsdale, 2004; King and Weber, 2007) and it is carried out by Actinobacteria, Proteobacteria and taxa from Bacteroidetes and *Chloroflexi* (Martín-Cuadrado *et al*., 2009) that were also detected in the Mediterranean deep sea metagenomes (Martín-Cuadrado *et al*., 2007). The ubiquitous distribution of the oxidation of carbon monoxide (CO-oxidation) by cox genes (K03518: *coxS* and K03519: *cosM*) was notorious, detected in nearly all samples and in high abundance (average 20% of the cells) (**Fig. 3**) and mostly associated to free-living microorganisms (Wilcoxon test p-value < 0.005) **(Fig.4)** pointing to CO-oxidation as an important energy supplement for heterotrophs in the deep ocean.

Anaerobic metabolisms, such as dissimilatory nitrate reduction (K00362, *nirB* -K00363, *nirD*), sulfate reduction (K00394, *aprA*-K00395, *aprB*), methanogenesis (K00320, *mer*; K14083, *mttB*) and denitrification (K00368, *nirK*; K02305, *norC*) appeared in between 60% and 100% of the samples (**Fig. 3, Table S3**). The prevalence of such metabolisms in well-oxygenated waters might be explained by the formation of microenvironments inside organic aggregates or particles, where concentrated respiration results in local O_2_ exhaustion. The dissimilatory nitrate reduction pathways and denitrification were both enriched in the particle-attached fraction (**Fig. 4**). While the dissimilatory nitrate reduction was present in most of the samples and in abundances that reached up to 34% of the microbial cells (**Fig. 3**), denitrification was less abundant (between 2-8% cells) despite being widely distributed. Indeed, different functional modules related to methanogenesis were found in about 17% of the microbial cells (**Fig. 3**) and significantly (Wilcoxon test p-value < 0.005) enriched in the free-living microorganisms (**Fig. 4**). Finally, the dissimilatory sulfate reduction genes were found in most of the FL and PA microbial communities and in 5-13 % of the microbial cells (**Fig. 3**). Our results not only highlight the distribution of the main biogeochemical processes in the bathypelagic oceans, some of them displaying a patchy distribution, but also disentangle specific metabolic pathways differentially associated to either the free-living or particle-attached microorganisms in the bathypelagic ocean, where these communities, therefore, contribute with different functions.

### Metabolic versatility and prevalence of mixotrophy in the deep ocean

The co-assembly of the 58 bathypelagic metagenomes allowed the reconstruction of 76 non-redundant and manually-curated metagenomic assembled genomes and 41 were taxonomically assigned to phylum or class by two independent approaches (see methods, **Fig. 1D**). This number is rather low compared to other initiatives from the photic oceans, likely reflecting the lower sequencing effort and lower diversity in the bathypelagic ocean. Our MAGs contained a total of 81,688,051 bp. 72 MAGs were taxonomically assigned to bacteria, and one was assigned to a fungus from genus *Tilletiaria* that was previously detected by 18S pyrotagging in the bathypelagic of several stations (**Fig. S1;** Pernice *et al*., 2016). The three remaining MAGs correspond to potential viruses of unknown taxonomy with genome sizes ranging from 127 to 215 Kb. *Alphaproteobacteria* (n=22) followed by *Gammaproteobacteria* (n=18), *Flavobacteriia* (n=7) and *Actinobacteria* (n=6) were the most abundant bacterial classes within our bathypelagic MAGs (**Fig. 1D**). A 14% of the MAGs (11 out of 76) represent potentially new phyla (**Table S4**). MAG completion estimations based on domain-specific single-copy core genes (Parks *et al*., 2015) ranged from 4.2 to 97.8% (**Table S4**), and 18 MAGs fitted the conditions of either high or medium-quality standards promoted for uncultured genomes at least with >50% genome completion and less that 10% contamination (Bowers *et al*., 2017; Konstantinidis *et al*., 2017) (**Table S4**). The 72 estimated bacterial MAGs sizes ranged from less than 1 Mbp up to 11 Mbp with an average estimated genome size of 3.62 Mbp **(Table S3).** These bathypelagic genomes are slightly larger in size than those estimated for bacteria and archaea in the 93 genomes from the photic ocean (surface and the deep chlorophyll maximum layer) obtained from the *Tara* Oceans project, which had 2.26 Mbp on average (Delmont *et al*., 2018). The larger genome size of the bathypelagic MAGs is in agreement with previous findings (Konstantinidis *et al*., 2009) and likely reflect higher metabolic versatility in resource utilization or complex biotic interactions (Konstantinidis, K. T. and Tiedje, 2004; Ranea, *et al*., 2005).

The bathypelagic MAGs represent a unique dataset to investigate the distribution and functional capacity of deep ocean microorganisms with the possibility to accurately link metabolic pathways to taxonomy (**Fig. 5).** Almost one fourth (24%) of the bathypelagic MAGs included mixotrophy, with the potential to perform both chemolithoautotrophy and heterotrophy. The mixotrophic MAGs occurred in > 75% of the samples pointing out the ubiquitous role of mixotrophy in the deep ocean (**Fig. 5**). We detected 10 out of the 76 MAGs (13%) that included two pathways for inorganic carbon fixation, the Calvin Cycle and the 3-Hydroxypropionate bicycle (**Fig. 5**). Out of the three MAGs with key genes of the Calvin Cycle, MAG6, related to Alphaproteobacteria (77.79% of estimated genome completeness) had the rubisco (*rbc*L) and the phosphoribulokinase (PRK) genes and half of the Calvin Cycle pathway genes (**Table S5**), but displayed also the complete SOX system for thiosulphate oxidation that provides energy for fixing CO_2_. Additionally, MAG12 (55% of genome completeness), related to *Methylophaga*, harbored also the phosphoribulokinase and 30% of the SOX system for thiosulphate oxidation, pointing to a mixotrophy lifestyle as previously described for *Methylococcus capsulatus* (Taylor *et al*., 1980; Baxter *et al*., 2002). These results provide evidence of great versatility and a prevalence of mixotrophy within the deep ocean microbiome.

**Figure 5.**
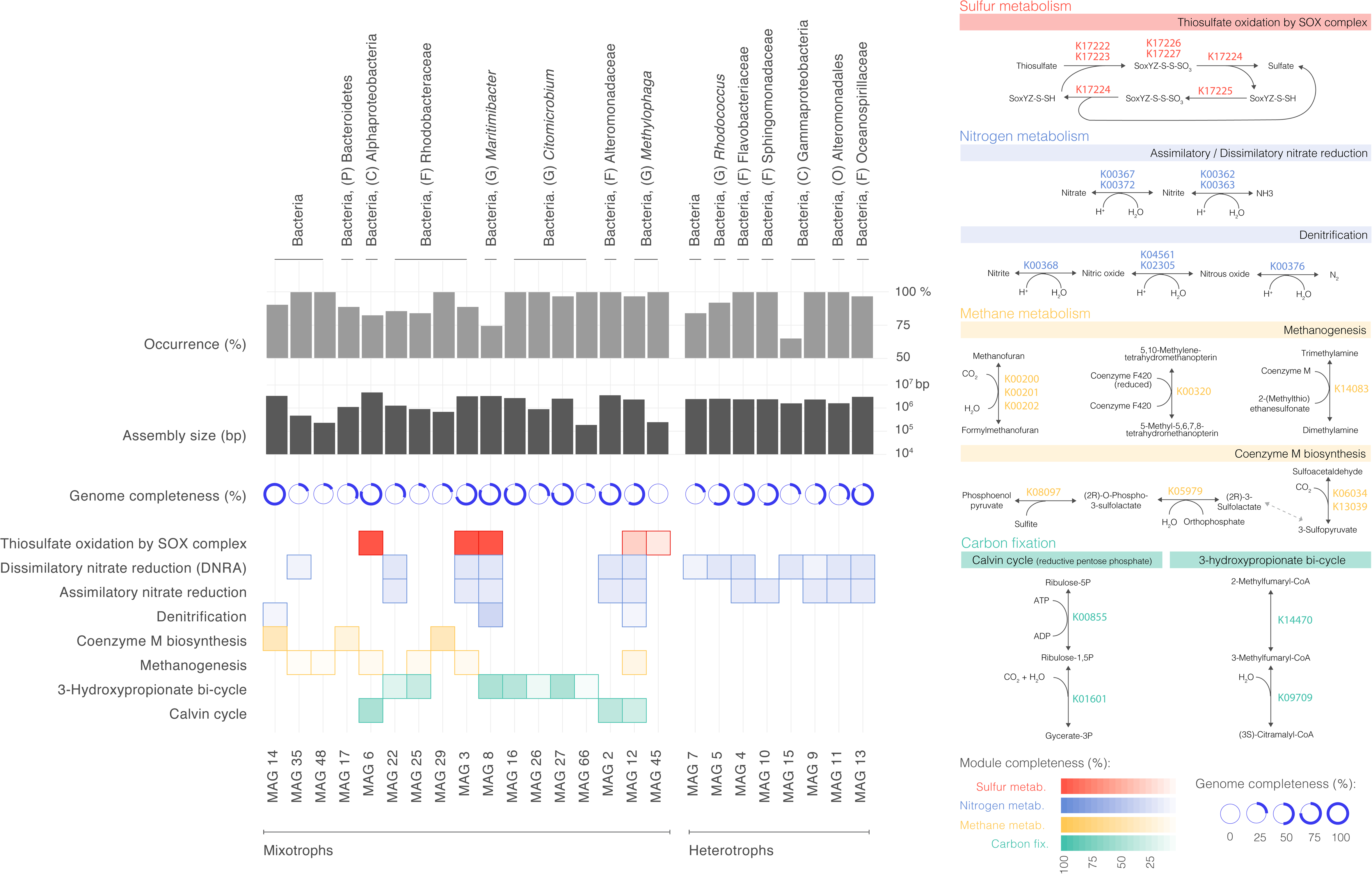
Metabolic pathways and potential mixotrophy in selected metagenome assembled genomes (MAGs) from the bathypelagic ocean. A total of 25 MAGs are highlighted based on the presence of metabolic pathways with potential for mixotrophy and some selected heterotrophs with assimilatory and dissimilatory nitrate reduction (DNRA). A total of 28 gene markers were found. Their taxonomic assignation to the maximum possible resolution is shown on the top, followed by the occurrence of each MAG in samples of the Malaspina Deep-Sea Gene Collection. The genome completeness (%) is also shown for each MAG. On the right, metabolic pathways involved in inorganic carbon fixation (green), sulfur (red), nitrogen (blue) and methane metabolism (orange) are shown as well as the KOs that unambiguously participate in these pathways. At the bottom, module completeness percentage for each pathway is coded by color intensity.

The most unexpected result was the prevalence of the 3-Hydroxypropionate bi-cycle in both the Malaspina Dee-Sea Gene Collection (**Fig. 3**) and the bathypelagic MAGs (**Fig. 5)**, which represents a pathway previously described with a narrow distribution within members of *Chloroflexaceae* (Zarzycki *et al*., 2009). Genes of this pathway have recently been detected within the uncultured SAR202 deep ocean clade genomes related to *Chloroflexi*, although the role of the 3-Hydroxypropionate bi-cycle in this clade has been linked to the assimilation of intermediate metabolisms produced by the degradation of recalcitrant dissolved organic matter (DOM) rather than for fixing CO_2_ (Landry *et al*., 2017). Thus, this clade displays the capacity to metabolize multiple organosulfur compounds and appears to be a sulfite oxidizer (Mehrshad *et al*., 2018). Seven MAGs (9% of the total) have at least two genes of the 3-Hydroxypropionate bi-cycle (K09709, *meh*; 3-methylfumaryl-CoA hydratase and K14470, *mct*; 2-methylfumaryl-CoA isomerase). None of them were assigned to the SAR202 clade, but instead were all related to Alphaproteobacteria, three taxonomically linked to *Rhodobacteraceae* and four to *Citromicrobium* (**Fig. 5**). In particular, MAG8, closely related to *Maritimibacter* (80% of estimated genome completeness) had a 48% pathway completeness. In these MAGs the key enzyme Malyl-CoA lyase (mcl, K08691) was also detected. Energy could be supplied by sulfur oxidation with the Sox system (pathway 100% complete). (**Fig. 5**). We detected the 3-Hydroxypropionate bi-cycle in the four MAGs associated with *Citromicrobium* (MAG16, MAG26, MAG27, MAG66). Two of them, with >75% of genome completeness, included at least 43% of this pathway. *Citromicrobium* MAGs were abundant in different stations and oceans (**Fig. 5**, see occurrence) and two of them contained the gene for the light-harvesting pigment bacteriochlorophyll (*puf*M) that provides capability for anoxygenic photosynthesis. The 3-hydroxypropionate autotrophic pathway was also found in other anoxygenic phototrophic bacteria related to the filamentous *Roseiflexus* spp., that belong to phototrophic Chloroflexi inhabiting microbial mats of hot springs (Klatt *et al*., 2007). Our *Citromicrobium* MAGs were closely related to *Citromicrobium bathyomarinum* isolated from black smoker plume waters of the Juan de Fuca Ridge at 2000 m deep in the Pacific Ocean (Yurkov and Beatty 1998). Their wide distribution across the bathypelagic ocean is consistent with the finding of these organisms in hydrothermal vent environments, where the high temperature could generate sufficient thermal radiation light to support anoxygenic photosynthesis in both mixotrophic or obligated organisms (Xiong *et al*., 2000; White *et al*., 2002; Beatty *et al*., 2005, Raven, J.A., 2009).

Although we lack experimental evidence that these MAGs can indeed perform inorganic carbon fixation, our results reveal their potential genetic capacity and motivate future experimental designs to unveil the ecological relevance of the 3-Hydroxypropionate bi-cycle in the deep ocean. In all these potentially chemolithoautotrophic MAGs, genes for central carbohydrate metabolisms such as the TCA cycle, glycolysis/gluconeogenesis or the pentose phosphate pathways were inferred with different degrees of completeness, that ranged from 8 to 46% (**Table S5**), revealing that mixotrophy with the ability to switch between autotrophic and heterotrophic metabolisms is prevalent in the bathypelagic ocean reflecting the metabolic versatility within the deep ocean microbiome. These findings, together with the enrichment of different metabolic pathways associated to either particle-attached and free-living microorganisms, expand our view of the metabolic seascape of the deep ocean microbiome.

## CONCLUSIONS

Our global metagenomic assessment of the deep ocean microbiome generating the Malaspina Deep-Sea Gene Collection and the bathypelagic MAGs datasets uncovers potential novel phyla and a description the main biogeochemical processes of the deep ocean microbiome of the tropical and subtropical bathypelagic ocean. The results provide evidence for a metabolic differentiation, reflected in contrasted functional gene repertories, between the particle-attached and the free-living microbial assemblages and provides evidence of a strategy towards metabolic versatility within the deep ocean. The widespread distribution of different autotrophic pathways in the deep ocean, specially the Calvin Cycle and the 3-Hydroxypropionate bi-cycle, and the prevalence of alternative energy sources such as CO-oxidation, sulfur oxidation and H_2_-oxidation, provides support for the role of a multitude of autotrophic processes in subsidizing the heterotrophic metabolism supported by export flux from the photic layer. This allows respiratory demands in the deep-sea to exceed the limits imposed by the carbon supply from the photic layer. The significance of mixotrophy and autotrophy in the deep-sea microbiome suggested here would need to be incorporated into current depictions of the deep ocean as relatively modest but necessary contributors to the global carbon budget.

## Acknowledgements

This work was funded by the Spanish Ministry of Economy and Competitiveness (MINECO) through the Consolider-Ingenio program (Malaspina 2010 Expedition, ref. CSD2008-00077). The sequencing of 58 bathypelagic metagenomes was done by the U.S. Department of Energy Joint Genome Institute, supported by the Office of Science of the U.S. Department of Energy under Contract No. DE-AC02 05CH11231 to SGA. Additional funding was provided by the project MAGGY (CTM2017-87736-R) to SGA from the Spanish Ministry of Economy and Competitiveness, Grup de Recerca 2017SGR/1568 from Generalitat de Catalunya, and King Abdullah University of Science and Technology (KAUST) under contract OSR #3362. High-Performance computing analyses were run at the Marine Bioinformatics Service (MARBITS) of the Institut de Ciències del Mar (ICM-CSIC), Barcelona Supercomputing Center (Grant BCV-2013-2-0001) and KAUST’s Ibex HPC. We thank the R/V Hesperides crew, the chief scientists in Malaspina legs, and all project participants for their help in making this project possible. We thank Shook Studio for assistance with figure design and execution.

