## Supplementary Figures for "Metabolic Architecture of the Deep Ocean Microbiome"

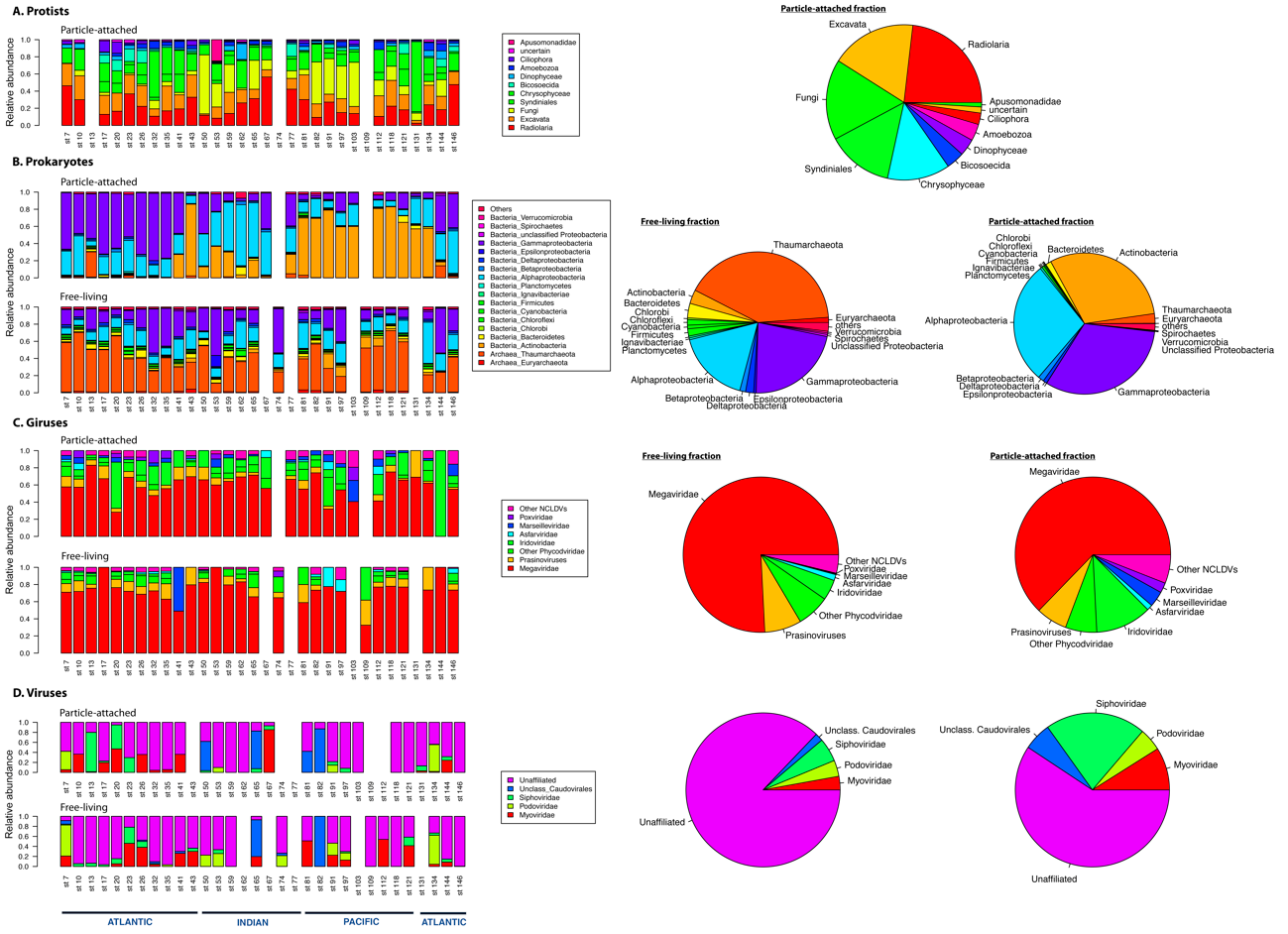

**Figure S1. Relative abundance of protists, prokaryotes, giruses, and viruses in the bathypelagic ocean.** A) Protists (small eukaryotes) only in the 0.8-20 µm, B) Prokaryotes (Bacteria and Archaea), C) Giruses and D) Viruses (prophage genes in metagenomes) from the Malaspina bathypelagic metagenomes. Different gene markers or strategies were used to assess the diversity and compute abundances from the metagenomes: i) for protist the 18S miTags approach was used, ii) for prokaryotes we used clade-specific marker genes from 3,000 reference genomes of Archaea and Bacteria to generate taxonomic abundance profiles in each sample, iii) the marker gene of the Nucleo-cytoplasmic Large DNA Viruses (NCLDV) major capsid was used for giruses and iv) the marker gene of the large subunit of the Terminase (TerL) was used for viruses. The left panel presents the relative abundance of each group per station split into the two size fractions (particle attached, PA, and Free-living, FL). The X-axis shows station (St) and oceans (Atlantic, Indian and Pacific). The right panel shows in pie charts the relative abundances of picoeukaryotes, prokaryotes, giruses and viruses as averages of all the stations.

A

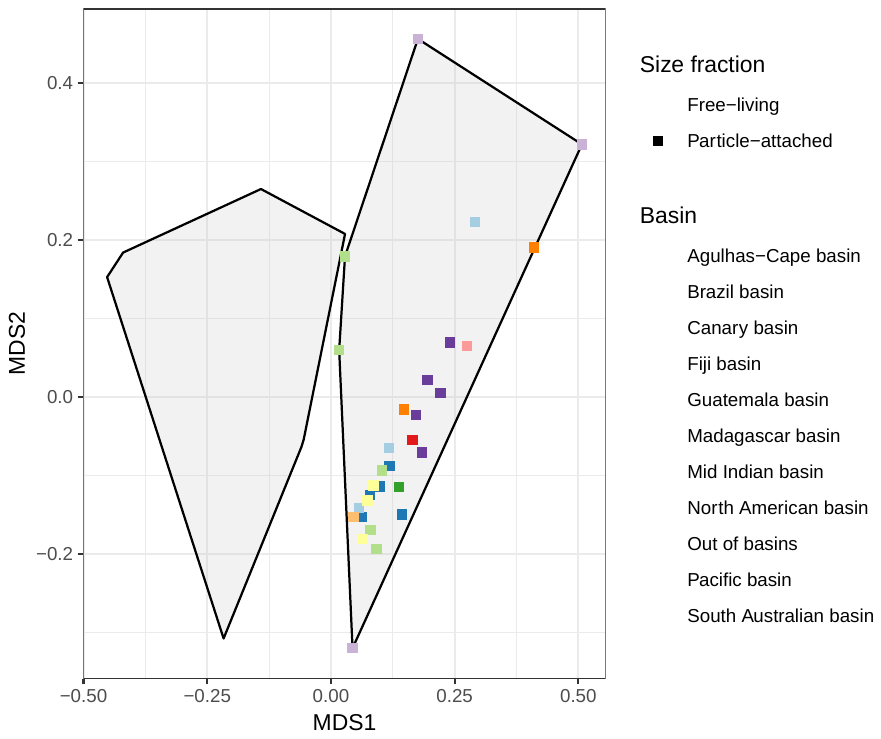

B

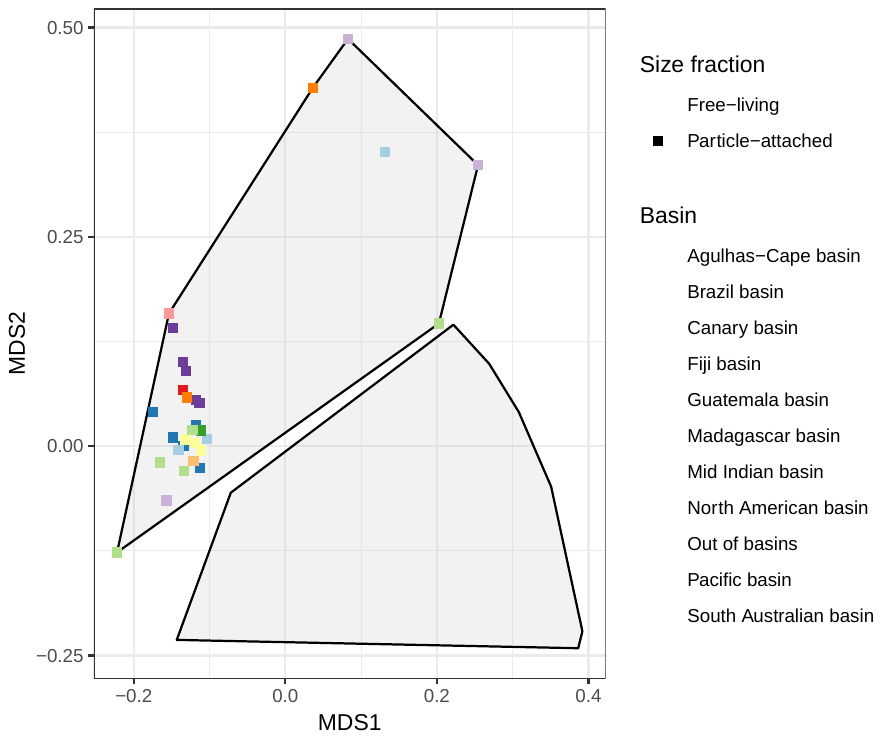

C

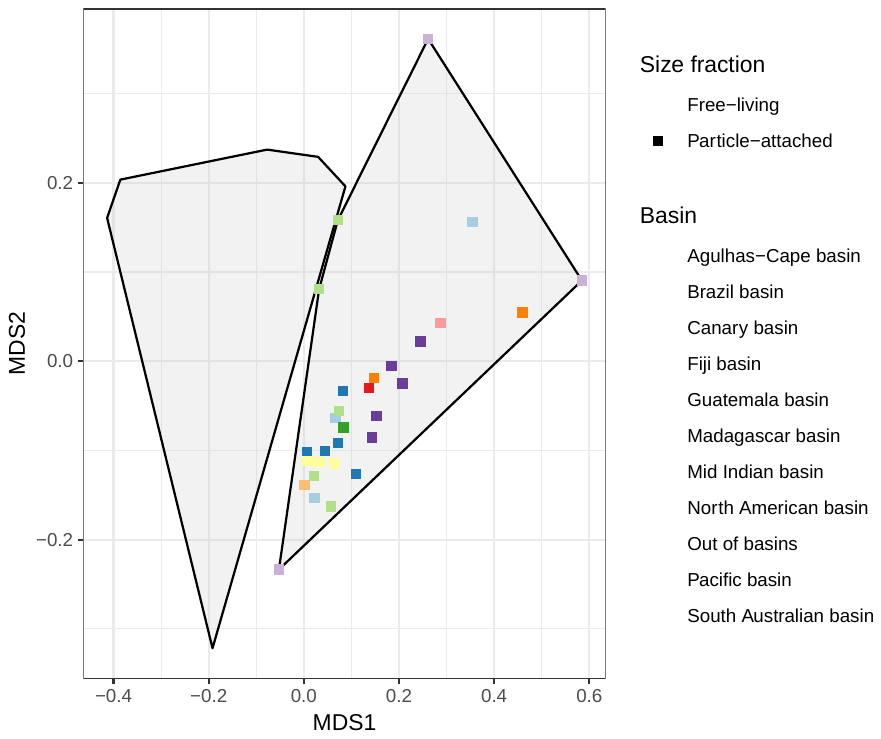

**Figure S2. Functional structure of the microbial deep ocean communities.** Bray Curtis distances among sites using non-metric multidimensional scaling (NMDS) ordination plots based on the functional abundance tables constructed with A) Protein families (Pfams) and B) Enzyme Commission numbers (ECs) and Cluster of Orthologous groups (COGs).
