## Supplementary Material and Methods for "Metabolic Architecture of the Deep Ocean Microbiome"

**Title:**

**Material and Methods:**

- **Sample collection and DNA extraction**
- **Library preparation and sequencing**
- **Data acquisition**
- **Generation of the Malaspina Deep-Sea Gene Collection (MDSGC)**
- **Taxonomic assignation of the metagenomic reads: from viruses to protists**
- **Marker enzymes from energy metabolisms**
- **Statistical analyses**
- **Co-assembly of bathypelagic metagenomes**
- **Generation of metagenomic assembled genomes (MAGs) from the Malaspina metagenomic dataset.**
- **Metabolic prediction of the MAGs.**
- **Estimation of module pathway completeness within the MAGs dataset**

**Sample collection and DNA extraction**

A total of 58 water samples were taken during the Malaspina 2010 expedition (http://www.expedicionmalaspina.es) corresponding to 32 different sampling stations globally distributed across the world’s oceans (**Figure 1A**). We focused on the samples collected at the depth of 4000 m, although a few samples were taken at shallower depths, all within the bathypelagic realm (average depth: 3731 m ± 495; standard deviation) in the tropical and subtropical oceans. Two different size fractions were analyzed in each station representing the free-living (FL, 0.2– 0.8 μm) and particle-attached (PA, 0.8– 20 μm) prokaryotic communities (Crump et al. 1999; Ghiglione et al. 2009; Allen et al. 2012). This size fractionation separated two different microbial community assemblages, mostly composed by particle-attached prokaryotes (PA; 0.8–20 µm) and free living prokaryotes in the surrounding water (FL; 0.2–0.8 µm) as previously shown (Salazar et al. 2015a, 2015b, Mestre et al. 2018). In addition, the PA assemblage also included microbial eukaryotes and their putative symbionts and the FL assemblage included some viruses.

For each sample 120 l of seawater were sequentially filtered through a 200 and a 20 μm mesh to remove large plankton. Further filtering was done by pumping water serially through 142-mm polycarbonate membrane filters of 0.8 μm (Merk Millipore, Darmstadt, Germany, Isopore polycarbonate) and 0.2 μm (Merck Millipore, Express Plus) pore size with a peristaltic pump (Masterflex, EW-77410-10). The filters were then flash-frozen in liquid N_2_ and stored at -80 °C until DNA extraction for whole community high-throughput shotgun sequencing.

The filters for metagenomic sequencing were cut in small pieces with sterile razor blades and half of each filter was used for DNA extractions, which were performed using the standard phenol-chloroform protocol with slight modifications (Logares et al*.* 2014). Details regarding the DNA extraction have been presented before (Salazar et al*.* 2015).

**Library preparation and sequencing**

Plate-based DNA library preparation for Illumina sequencing was performed on the PerkinElmer Sciclone NGS robotic liquid handling system using Kapa Biosystems’ library preparation kit. A total of 200 ng of sample DNA was sheared to 270 bp using a Covaris LE220 focused-ultrasonicator. The sheared DNA fragments were size selected by double-SPRI and then the selected fragments were end-repaired, A-tailed, and ligated with Illumina compatible sequencing adaptors from IDT containing a unique molecular index barcode for each sample library. The prepared libraries were then quantified using KAPA Biosystem’s next-generation sequencing library qPCR kit and run on a Roche LightCycler 480 real-time PCR instrument. The quantified libraries were then multiplexed into pools of 8 or 12 libraries each, and the pool was then prepared for sequencing in the DOE's Joint Genome Institute (JGI) on the Illumina HiSeq2000 sequencing platform utilizing a TruSeq paired-end cluster kit, v3, following a 2x150 indexed run recipe, and Illumina’s cBot instrument to generate a clustered flowcell for sequencing.

**Data acquisition**

Raw and clean sequences, were obtained from DOE's JGI Integrated Microbial Genomes and Microbiomes (IMG/MER; NCBI accession numbers available in **Table SM1**), as well as several analyzed data (Metagenome Annotation Standard Operating Procedure for IMG, Nov 2012) available at JGI proposal ID 300784). Functional abundance tables containing the number of reads in each sample for every functional category within four different functional annotations: Cluster of Orthologous Genes (COG), Clusters of KEGG orthologous genes (KOs), Protein families (Pfam) and Enzyme Comission classification (EC). In all cases, abundance tables were downloaded directly from IMG repository (accessions number in **Table 1SM**) using the “estimated gene copies” option, i.e. the abundance tables represented the read counts for every annotated function in every sample either coming from unassembled or assembled gene data (multiplying by read depth in the latter case).

Additional metadata were collected during the expedition including environmental variables (salinity, potential temperature and oxygen concentration), sampling station coordinates (latitude, longitude and depth) and auxiliary data for every sample (filter size, ocean basin and water mass) (see **Table 1SM**).

**Generation of the Malaspina Deep-Sea Gene Collection (MDSGC)**

All 3,872,410 predicted coding sequences larger than 100 bp from each assembled metagenome were pooled and clustered at 95% of sequence similarity and 90% of sequence overlap of the smaller sequence using cd-hit-est (v.4.6; Li and Godzik 2006) using the following options: -c 0.95 -T 0 -M 0 -G 0 -aS 0.9 -g 1 -r 1 -d 0 to obtain 1,115,269 non-redundant gene clusters (from now on referred simply as genes).

In order to explore the novelty of the MDSGC, we clustered it with the 40,154,822 non-redundant sequences from the *Tara* Oceans Microbial Reference Gene Catalog (OM-RGC; Sunagawa et al. 2015) using cd-hit-est-2d (v.4.6; Li and Godzik 2006) using the following options: -c 0.95 -T 24 -M 74000 -G 0 -aS 0.9 -g 1 -r 1 -d 0 to obtain a final catalog of 40,812,291 genes.

**Taxonomic assignation of protist, prokaryotes and viruses from metagenomic reads**

***Protist***

18S miTags were extracted from the metagenomes and subsequently analyzed following Logares et al., 2014. These miTags were mapped at 97% similarity using Uclust (Edgar,RC (2010) to the PR2 database (Guillou et al. 2013) that was pre-clustered at 97% similarity.

***Prokaryotes***

The software MetaPhlAn (Segata et al., 2012) was used to align cleaned and trimmed metagenomic reads to a catalog of clade-specific marker genes identified from 3,000 reference genomes of Archaea and Bacteria, and to generate taxonomic abundance profiles in each sample.

***Nucleo-cytoplasmic large DNA viruses (NCLDV)***

Nucleo-cytoplasmic large DNA viruses (NCLDV) marker genes, including major capsid proteins and DNA polymerases, were detected in the 58 Malaspina deep metagenomics samples with the use of previously described procedures (Hingamp et al., 2013) using NCVOG (Yutin et al, 2009) and PSI-BLAST (E-value < 1e-3) (Altschul et al. 1997). For the detection of virophage sequences, we first screened the metagenomic sequences by the proteome sequences of three virophages (Sputnik, Mavirus, OLV) by BLAST (E-value<0.001). Then the metagenomic hits were PSI-BLAST (E-value < 1e-3) against UniRef100 (Suzek et al. 2015). As a result, 365 metagenomic peptides best hit to virophage sequences (34 distinct genes).

***Viral signal analysis***

The marker gene TerL (large subunit of the Terminase) was used to assess the diversity of bacterial and archaeal viruses in the deep-sea microbial metagenomes. TerL genes were identified in the proteins predicted from the 58 metagenomes through hits to the PFAM domains PF04466 (Terminase_3), PF03237 (Terminase_6), PF03354 (Terminase_1) and PF05876 (Terminase_GpA). Genes shorter than 100 bp were discarded, leading to a dataset of 485 TerL genes. First, a family-level affiliation (i.e. *Myoviridae*, *Podoviridae*, or *Siphoviridae*) of these sequences was obtained from a best blastp hit to the RefseqVirus database (threshold of 50 on bit score, 0.001 on e-value, and 50 on % of amino acid identity). For each sample, a viral community composition was then calculated based on this family-level affiliation and the normalized coverage of the contig (i.e. contig coverage divided by contig length and sequencing depth of the sample, as in Brum et al., 2015). Next, these 485 “deep-sea” TerL genes were clustered with all TerL from the RefseqVirus database (n=899, v72, 09-2015), from “environmental phages” in Genbank (n=456, downloaded on 07-2015), from the VirSorter Curated Dataset (n=6,600, Roux et al., 2015), and from the Global Ocean Virome Dataset (n=2,674, sequences from 91 Epi- and Mesopelagic viromes from Tara Oceans and Malaspina expedition, Roux et al., 2016) at 98% of nucleotide identity (threshold most consistent with genome-based population definition, i.e. ≥80% of genes shared at ≥95% average nucleotide identity, when tested on complete genomes from RefseqVirus), leading to 5,701 OTUs. The 485 TerL genes from Malaspina were distributed across 303 OTUs, including 300 unique to deep-sea samples (i.e. containing only Malaspina TerL sequences).

**Marker enzymes from energy metabolisms**

Metabolisms with a key role in the main biogeochemical cycles in the deep ocean were studied with additional detail by choosing specific enzymes for nitrogen, sulfur, methane, hydrogen and carbon fixation metabolic pathways. Marker enzymes for different pathways were defined by exploring the corresponding Kyoto Encyclopedia of Genes and Genomes (KEGG) pathway maps (KEGG release 76.0; Kaneisha and Goto, 2000). Enzymes participating only in reactions within each map were defined as marker enzymes for this metabolic map. When possible, marker enzymes were assigned to a specific module within a map (e.g. enzymes only participating in Denitrification module within the Nitrogen metabolism map). Corrected abundance estimation for marker enzymes was obtained by dividing the number of reads from the EC functional abundance table (without subsampling) by the number of reads assigned to the prokaryotic single-copy gene *recA* (selected as COG0468). A table selection of 69 KOs was built (**Table 2SM**) representing main key markers genes for different metabolic pathways relevant in the deep ocean. Of those a total of 40 KOs were found in the Malaspina Deep-Sea Gene Collection (58%).

**Statistical analyses**

For every functional abundance table, a subsampled equivalent table was constructed in order to avoid biases due to the varying sequencing depth between samples. Subsampling was performed by generating a randomly rarefied table without replacement from the original one with “rrarefy” function in *vegan* package within R software (R Core Team 2012).

**Non-metric multidimensional scaling (NMDS)**

Non-metric multidimensional scaling (NMDS) with stable solution from random starts was performed for ordination of samples based on functional similarity using the different functional abundance tables (see **Fig. 2** and **Fig. S2**). Bray-Curtis distance measure was used for the abundance tables. Wisconsin double standardization and square root transformation (Bray & Curtis 1957) were applied before performing the analysis. All NMDS analyses were performed with the subsampled version of each abundance table.

**Metagenomic assembled genomes from the bathypelagic ocean**

Metagenome Assembled Genomes (MAGs) were obtained by binning contigs larger than 100 kbp resulting from a co-assembly of all 58 metagenomes from the Malaspina expedition (Ray mèta parallel de novo genome assembler v2.2.0; Boisvert et al., 2012). The co-assembly allowed us to increase the sequence space and we obtained a total of 152,175 contigs larger than 2,000 bp. For the downstream analyses only those contigs larger than 100 Kbp were kept, accounting for a total of 448 contigs.

The abundance of each MAG was assessed by mapping competitively the reads from the 58 metagenomes against the contigs database using blastn (Nucleotide-Nucleotide BLAST 2.2.26+) using the same parameters as in Swan et al. (2013). The mapped metagenomic reads that aligned in less than 90% of its length were excluded for downstream analyses. Likewise, we kept only those metagenomic reads with an identity higher than 95% to a contig in the reference database and, in this case, the metagenomic read was assumed that belonged to the reference contig (Caro-Quintero et al., 2011). The number of reads recruited per contig was normalized by the contig size in all samples to avoid biases generated by differences in genome length. All samples were randomly subsampled to the minimum number of reads mapped in one sample. The normalized abundance of each contig along the samples was used to predict and build MAGs. Contigs were considered belonging to the same MAG if their abundance patterns were significantly correlated (*P* < 10^-5^, linear regression analysis), the slope of the linear regression were 1 (0.1) and the R^2^ of the adjustment were >0.9. The contigs binned with this method were confirmed with metaBAT (Kang et al. 2015). A total of 76 MAGs were predicted. The completeness of the MAGs was estimated using both the standalone version of fetchMG v1.0 (Kultima et al., 2012) and checkM v1.0.5 (Parks et al. 2014) to eventually keep the finest taxonomic rank shared between methods.

**Metabolic prediction of the MAGs.**

The Kyoto Encyclopedia of Genes and Genomes (KEGG release 76.0; Kaneisha and Goto, 2000), and particularly, the KEGG orthology (KO) database, was used to determine the energetic metabolism of the Malaspina Deep-Sea MAGs. A total of 69 marker genes from carbon fixation, methane, nitrogen, hydrogen and sulfur metabolisms were selected (**Table 2SM**) and their presence was explored along the 76 MAGs. A total of 28 KOs was found in the MAGs dataset.

**Estimation of module pathway completeness within the MAGs dataset**

Module completeness per MAG was estimated calculating the percentage of KOs belonging to each module over its total KO number (KEGG release 76.0).
